# In vitro to in vivo evidence for chemical disruption of glucocorticoid receptor signaling

**DOI:** 10.1101/2025.03.07.642104

**Authors:** Maeve T. Morris, Jordan L. Pascoe, Jonathan T. Busada

**Author notes:** **Corresponding Author:** Jonathan T. Busada, WVU School of Medicine, Microbiology, Immunology and Cell Biology, 64 Medical Center Drive, P.O. Box 9177, Morgantown, WV 26506. **Author Contributions:** M.T.M. and J.T.B. planned the study. M.T.M. and J.L.P. performed all experiments. M.T.M. and J.T.B. analyzed all data and drafted the manuscript. All authors reviewed and approved the final manuscript.

## Abstract

Glucocorticoids are steroid hormones that regulate stress homeostasis, metabolism, and inflammatory responses. Dysregulation of the glucocorticoid receptor (GR) is linked to diseases such as obesity, mood disorders, and immune dysfunction. Endocrine-disrupting chemicals (EDCs) are widespread environmental contaminants known to interfere with hormone signaling, but their impact on glucocorticoid signaling remains unclear. While several GR-disrupting compounds have been identified *in vitro*, their *in vivo* effects remain largely unknown. In this study, we identified the agricultural agents dichlorodiphenyltrichloroethane (DDT) and ziram as GR-disruptors *in vitro. In vivo*, corticosterone co-treatment with DDT or the GR antagonist RU-486 inhibited the expression of classic GR-regulated transcripts in the liver. Furthermore, chronic exposure to DDT or RU-486 significantly reduced circulating B and T lymphocyte populations, respectively. These findings underscore the need to translate *in vitro* discoveries into *in vivo* models to assess the clinical relevance of GR-disrupting compounds. Moreover, they highlight the potential for xenobiotic-induced GR disruption to impair metabolic and immune homeostasis, potentially increasing disease susceptibility.

## Introduction

Glucocorticoids are steroid hormones produced by the adrenal glands at the end of the hypothalamic-pituitary-adrenal (HPA) axis that play an essential role in maintaining homeostasis, regulating metabolism, immune function, and the body’s response to stress [1, 2]. They exert their physiological effects through the glucocorticoid receptor (GR), a ligand-dependent transcription factor that mediates changes in gene expression upon binding to glucocorticoid response elements (GREs) on the DNA [3]. These hormones are crucial for processes such as glucose metabolism and inflammation. Dysregulation of glucocorticoid signaling has been implicated in a wide range of health conditions, including metabolic disorders, immune dysfunction, cardiovascular diseases, and mood disorders, and may increase the susceptibility of developing neoplasia [4-6].

Endocrine-disrupting compounds (EDCs) are environmental chemicals that can interfere with hormone signaling and disrupt the normal functioning of the endocrine system [7]. Exposure to EDCs, such as pesticides, industrial chemicals, and plastics, is ubiquitous and persistent, posing a growing concern for public health. These compounds are known to interfere with various aspects of hormone signaling, including steroid hormones like estrogen, androgens, and glucocorticoids [8]. Despite the growing recognition of the effects of EDCs on the endocrine system, the specific mechanisms by which they disrupt glucocorticoid signaling and their role in disease remain poorly understood.

Several EDCs have been reported to affect GR signaling *in vitro*, but their impact *in vivo* remains poorly understood [9, 10]. Establishing *in vivo* testing guidelines is crucial for validating GR-disrupting compounds and assessing their long-term effects on glucocorticoid-regulated physiological processes. This study aims to identify potential EDCs, evaluate their effects on GR transactivation *in vitro*, and validate their GR-disruptive properties in mice. Through analyses of GR binding, nuclear translocation, and transactivation, we identified the agricultural agent’s dichlorodiphenyltrichloroethane (DDT) and ziram as endocrine disruptors. Further *in vivo* testing of DDT revealed that acute exposure blocked corticosterone activation of classical GR-regulated genes in the liver, while chronic exposure reduced the proportion of circulating B cells. These findings highlight both the importance and challenges of evaluating putative GR-targeting EDCs *in vivo*.

## Methods

### Animal Care and Treatment

All mouse studies were performed with approval by the Animal Care and Use Committee at West Virginia University. C57BL/6J mice were purchased from the Jackson Laboratories. All studies utilized female mice. All mice were administered standard chow and water *ab libitum* and maintained in a temperature- and humidity-controlled room with standard 12-hour light/dark cycles. All experiments, including intraperitoneal (IP) injections, adrenalectomy (ADX) surgeries, and chronic EDC exposures, were performed at 8 weeks of age. Following ADX, mice were sustained on 0.9% saline drinking water to maintain ionic homeostasis.

### Viability Assay

A549 cells were seeded into a 96-well plate at a density of 5,000 cells per well. Cells were treated with cortisol (Steraloids), RU486 (10µM) (Steraloids), or the putative EDC (10µM, 1µM, 0.1µM) for 24 hours, then the CCK-8 Cytotoxicity Assay was performed according to manufacturer protocol (ApeX Bio).

### Luciferase Assay

A549 cells were transfected using the TransIT-X2 Dynamic Delivery System (Mirus) with a plasmid containing a glucocorticoid-responsive luciferase (GRE2-LVC), where two glucocorticoid response elements were inserted upstream of the luc operon. Cells were pretreated with RU486 (10µM) or the chemical of interest (10µM, 1µM, 0.1µM) for 1 hour, then 10nM cortisol was added and incubated for 18 hours. The cells were then washed, and the Dual-Glo Luciferase assay (Promega) was used. The SpectraMax iD5 plate reader (Molecular Devices) was used to read luminescence.

### Lanthascreen Assay

Using Thermo Fisher Scientific’s SelectScreen Profile Service, RU486, o,p’-DDT (Chem Service Inc.), metolachlor (Sigma-Aldrich), and ziram (Sigma-Aldrich) were tested against the synthetic glucocorticoid dexamethasone for their GR binding ability using a Lanthascreen competitive binding assay.

### RNA Isolation from Cell Culture and qRT-PCR

A549 cells were seeded into a 12-well plate at a density of 75,000 cells/well. Cells were pretreated with a final concentration of RU486 (10µM), DDT (10µM, 1µM, 0.1µM), or ziram (10µM, 1µM, 0.1µM) for 1 hour. Then cortisol (10nM) was added, and cells were incubated for 6 hours. After 6 hours, A549 cells were lysed using the kit reagents, and the kit protocol was followed to isolate RNA (Omega Bio-Tek). Reverse transcription followed by qPCR was performed in the same reaction using the Universal Probes One-Step PCR kit (Bio-Rad Laboratories) and the TaqMan primers: PPIB (Hs00168719_m1), FKBP5(Hs01561006_m1), and PER1(Hs00242988_m1) (ThermoFisher).

### Glucocorticoid Translocation and Immunofluorescence Staining

A549 cells were seeded into a 35mm dish at a density of 150,000 cells. Cells were treated with cortisol (10nM) with or without the putative EDC (10µM, 1µM, 0.1µM) for 1.5 hours. Cells were fixed in 4% paraformaldehyde for 20 minutes, and then standard methods were used to perform immunofluorescence staining. Cells were incubated in Nr3c1 (D8H2) (Cell Signaling) overnight at 4°C. They were then incubated in secondary antibodies and Phalloidin-iFluor (abcam) for 1 hour at room temperature. Cells were mounted with Vectastain mounting media containing 4’,6-diamidino-2-phenylindole (Vector Laboratories). Images were obtained using a Zeiss 710 confocal laser-scanning microscope (Carl-Zeiss) and running Zen Black (Carl-Zeiss) imaging software.

### GR-EDC *In vivo* Exposures

For intraperitoneal (IP) injections, RU486 and DDT were dissolved in 100% ethanol, then resuspended in sterile 1X PBS to make a 1% ethanol solution. Mice were IP injected with 30mg/kg RU486 or 30mg/kg DDT. After 1 hour, mice were IP injected with 1mg/kg corticosterone. After 3 hours, the mice were euthanized. RU486 and DDT were prepared for drinking water exposure by dissolving the chemicals in ethanol and resuspending them in water to make a 1% ethanol solution. Mice received a final dose of 2mg/kg/day RU486 or 3mg/kg/day DDT through their drinking water. Fresh water was prepared twice each week.

### Corticosterone ELISA

Serum was collected by submandibular bleed at the end of the 2-week and 1-month exposures. Circulating corticosterone levels were measured by ELISA assay following the manufacturer’s protocol (Arbor Assays).

### RNA Isolation from Tissue and qRT-PCR

RNA was extracted in TRIzol (ThermoFisher) and precipitated from the aqueous phase using 1.5 volumes of 100% ethanol. The mixture was transferred to an RNA isolation column (Omega Bio-Tek), and the remaining steps were followed according to the manufacturer’s recommendations. Reverse transcription followed by qPCR was performed in the same reaction using the Universal Probes One-Step PCR kit (Bio-Rad Laboratories) and the TaqMan primers: *Ppib* (Mm00478295_m1), *Fkbp5* (Mm00487406_m1), *Sgk1* (Mm00441380_m1), *Tsc22d3* (Mm01304886_m1), *Nr3c1*(Mm00433832_m1), *Star* (Mm00441558_m1), and *Cyp11b1* (Mm01204952_m1) (all from Thermofisher).

### Flow Cytometry

Flow cytometry was done on blood collected by a cardiac stick. 100 µl of blood was extracted and incubated in ammonium-chloride-potassium (ACK) lysing buffer for 10 minutes on ice. Cells were stained with: CD3e (clone 17A2), CD4 (clone GK1.5), CD8a (clone 53-6.7), B220 (clone RA3-6B2), and CD45.2 (clone 104) (all from BioLegend), and 7AAD for 20 minutes on ice. Samples were analyzed on a Cytek Aurora spectral flow cytometer (Cytek Biosciences). Flow cytometry analysis was performed using Cytobank (Beckman Coulter).

### Statistics

All error bars are ± SD of the mean. The sample size for each experiment is indicated in the figure legends. Experiments were repeated a minimum of two times. A one-sample t-test was used when comparing one group. An unpaired t-test was used when comparing two groups. A one-way analysis of variance with the post hoc Tukey t-test was used when comparing three or more groups. Statistical analysis was performed using GraphPad Prism 10 software. Statistical significance was set at *p*≤0.05. Specific *P* values are listed in the figure legends.

## Results

### DDT and Ziram act as glucocorticoid receptor disruptors in vitro

Many studies have reported compounds that bind to the GR and disrupt transactivation *in vitro*. However, few have assessed putative EDCs in *in vivo* studies. We selected several putative GR-EDCs identified in the literature and screened them for their ability to disrupt GR transactivation (Supplementary Table 1). To establish a dose concentration, A549 cells were treated with the putative EDCs at 10µM, 1µM, and 0.1µM (Figure 1A). Treatment with the EDCs was based on the viability results. To screen the putative EDCs for their ability to disrupt GR transactivation, we used a GR-responsive luciferase assay in which A549 cells were transfected with a construct containing glucocorticoid response elements (GREs) coupled to a luciferase reporter [11]. Treatment with the known glucocorticoid cortisol induced high luciferase expression, but the addition of the known GR antagonist RU486 [12] significantly reduced cortisol-induced luciferase (Figure 1B). Surprisingly, the bulk of the tested compounds did not yield a convincing alteration of cortisol-induced luciferase activity (Supplementary Figure 1). However, treatment with the insecticides dichlorodiphenyltrichloroethane (DDT) and ziram and the herbicide metolachlor significantly reduced luciferase flux. Next, we treated A549 cells with cortisol with or without the putative GR-EDCs to determine their impact on the activation of the well-known GR-target genes *FKBP5* and *PER1* [13, 14]. Cortisol increased the expression of *FKBP5* and *PER1*, while RU486 significantly reduced gene expression. DDT, metolachlor, and ziram significantly reduced *FKBP5* and *PER1* expression (Figure 1C), suggesting they disrupt glucocorticoid-mediated gene transcription. Together, these results show DDT, metolachlor, and ziram disrupt GR transactivation.

**Figure 1.**
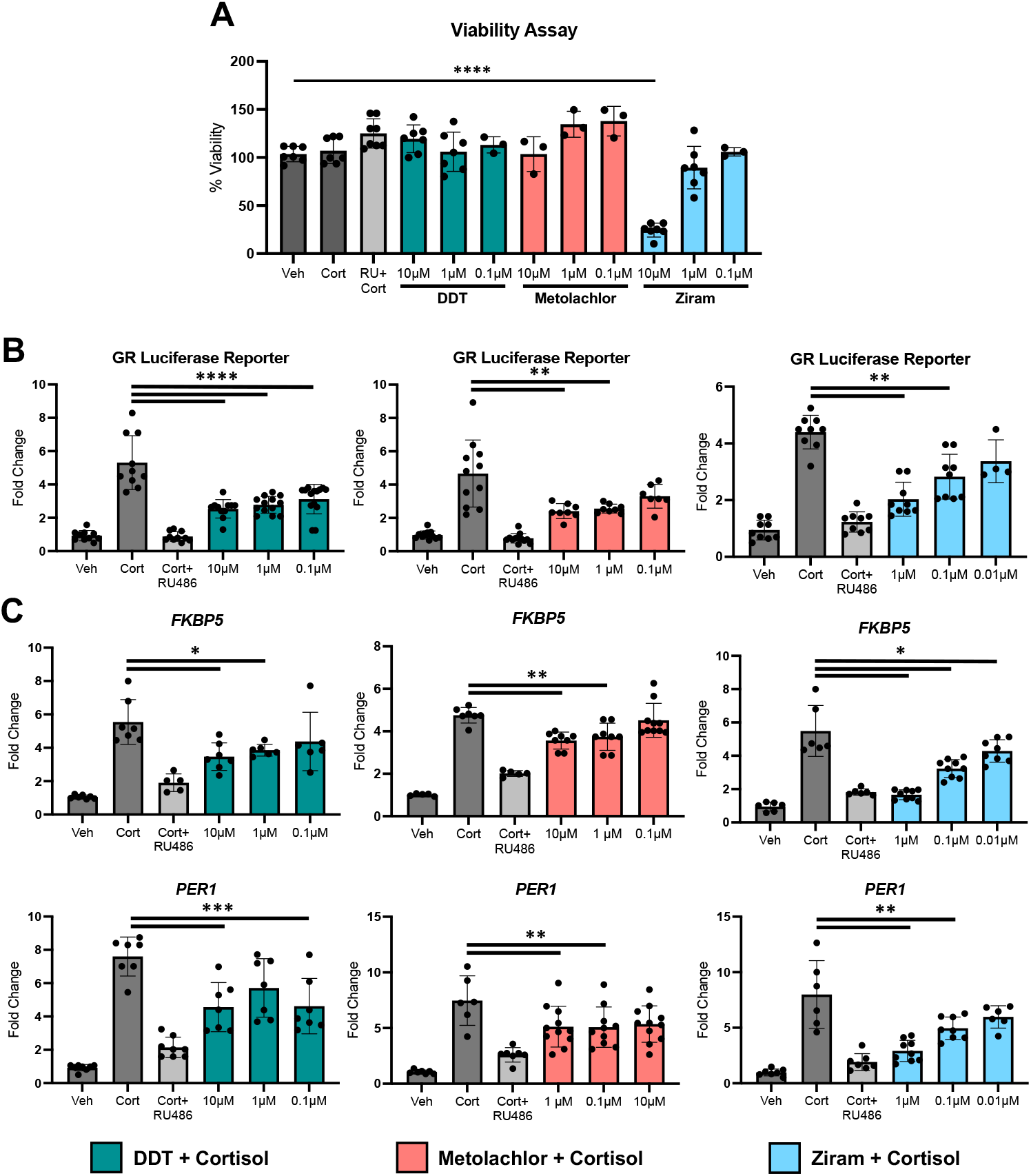
DDT, metolachlor, and ziram act as EDCs and reduce GR transactivation. (A) Viability of A549 cells was determined 24 hours after EDC treatment. (B) A549 cells were transfected with a glucocorticoid-responsive luciferase construct. Cells were pretreated with RU486 or the putative EDCs for 1 hour, and then cortisol was added for 18 hours. (C) A549 cells were pretreated with RU486 or the putative EDCs for 1 hour, then cortisol was added for 6 hours. qRT-PCR of GR target genes. n≥6. *p≤0.05, **p≤0.01, ***p≤0.001, and ****p≤0.0001.

Glucocorticoid signaling is a multistep process in which the ligand binds the GR, allowing nuclear translocation and binding GREs on the DNA [15]. Known EDCs have been shown to influence glucocorticoid activity at different stages. To evaluate DDT, metolachlor, and ziram for their ability to bind the GR, we utilized a Lanthascreen competitive binding assay. The putative EDCs were tested for their ability to outcompete the synthetic glucocorticoid dexamethasone for the GR. RU486 had a high affinity for the GR, with an IC50 of 1.32nM. DDT and ziram competitively bound the GR, but not until higher concentrations, with an IC50 of 2,720nM and 17,200nM, respectively). Metolachlor did not bind the GR at any of the tested concentrations and was, therefore, removed from the remaining studies (Figure 2A). These results indicate that DDT and ziram act as GR antagonists. Next, we further identified the disruptive effects of DDT and ziram using immunofluorescence staining of the GR, visualizing nuclear translocation. In unstimulated cells, the GR can be seen throughout the cellular cytoplasm and nucleus, while cortisol induced complete nuclear translocation. Interestingly, we found that DDT and ziram have different mechanisms of glucocorticoid disruption. DDT induced GR translocation to the nucleus in the absence of cortisol. Alternatively, ziram prevented GR translocation in the presence of cortisol (Figure 2B-C). This indicates two unique mechanisms by which DDT and ziram disrupt glucocorticoid signaling. Both can bind the GR, where DDT induces nuclear translocation but reduces transactivation. Ziram binds the GR, which holds it in the cytoplasm even in the presence of cortisol, leading to reduced transactivation. The *in vitro* studies demonstrate that DDT and ziram act as glucocorticoid disruptors.

**Figure 2.**
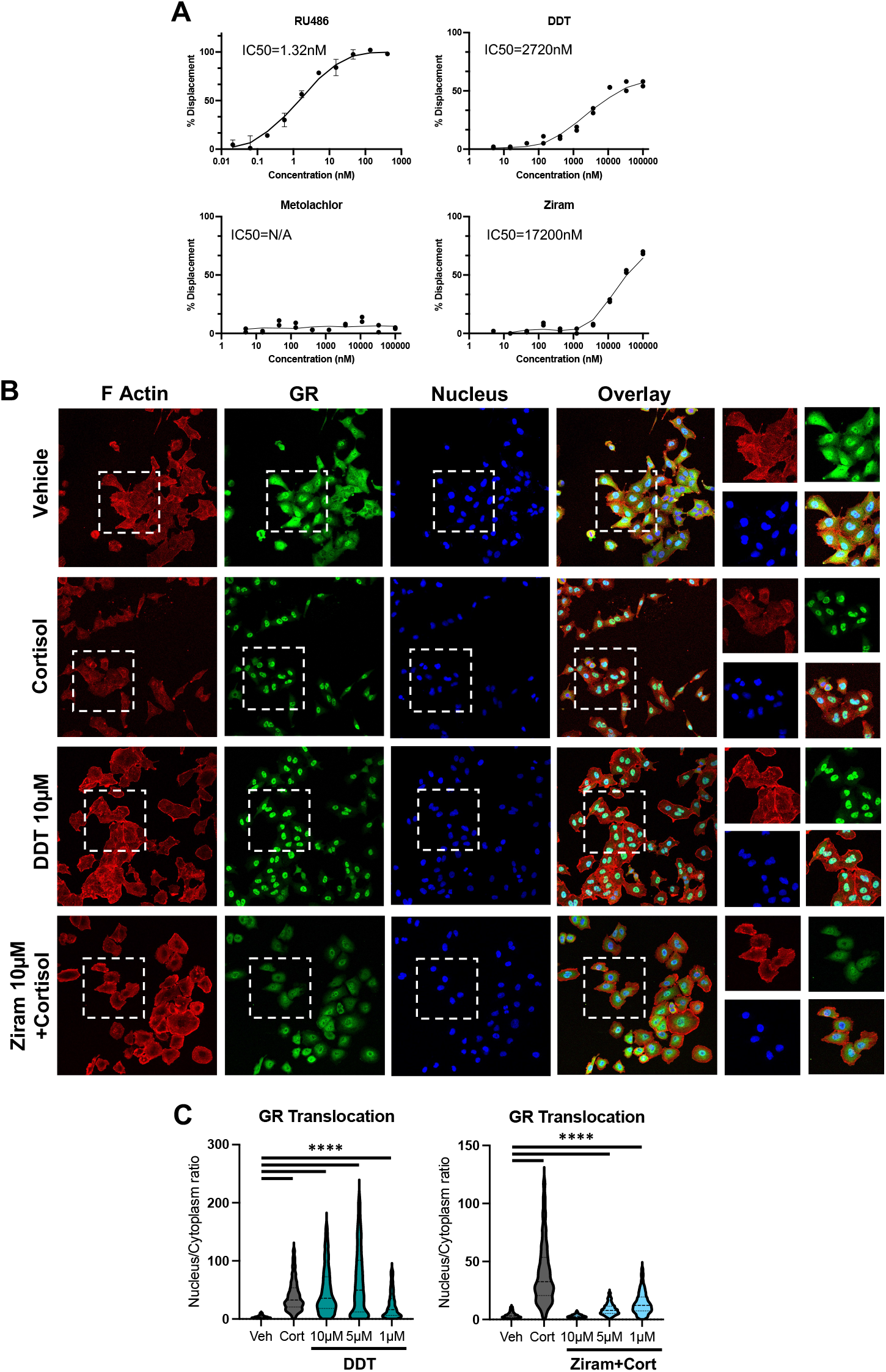
DDT and ziram act as GR antagonists. (A) Lanthascreen competitive binding assay assessing the putative EDC’s ability to outcompete dexamethasone for the GR. (B) Immunofluorescence imaging of GR translocation in A549 cells 1.5 hours after treatment with the EDCs. (C) The ratio of GR in the nucleus compared to the cytoplasm to quantify GR translocation. *p≤0.05, **p≤0.01, ***p≤0.001, and ****p≤0.0001.

### Acute exposure of DDT in vivo disrupts GR transactivation

We next sought to test the ability of the identified EDCs to disrupt GR *in vivo*. Few studies have used animal models to define the effects of glucocorticoid disruption. Endogenous glucocorticoids were removed from mice by performing bilateral ADX. Using an intraperitoneal (IP) injection, the mice were exposed to corticosterone with or without RU486 or DDT. After 3 hours, the liver was harvested, and qRT-PCR of GR-regulated genes was performed (Figure 3A). Ziram was removed from all *in vivo* studies due to its toxicity. Corticosterone increased the expression of *Fkbp5, Sgk1*, and *Tsc22d3* but not *Nr3c1*. RU486 significantly reduced corticosterone-induced expression of *Sgk1* and *Tsc22d3*. Interestingly, DDT reduced the expression of all tested genes (Figure 3B), indicating the disruptive effects of GR transactivation in acute exposures.

**Figure 3.**
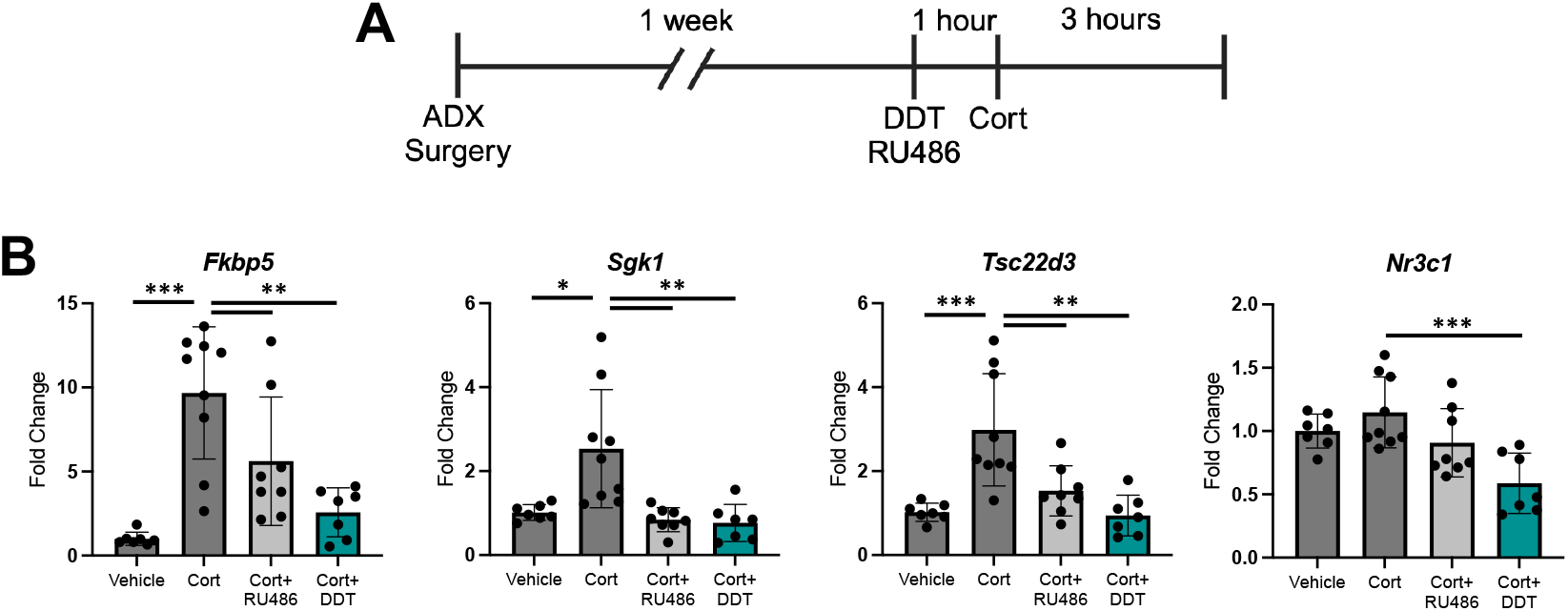
Acute in vivo exposure to DDT disrupts GR transactivation. (A) C57BL/6 mice underwent adrenalectomy surgery 1 week before EDC treatment. RU486 or DDT was delivered by IP injection 1 hour before injecting corticosterone. After 3 hours, the mice were euthanized, and the livers were harvested. (B) qRT-PCR of glucocorticoid responsive genes using RNA isolated from the liver. n≥7. *p≤0.05, **p≤0.01, ***p≤0.001, and ****p≤0.0001.

### Chronic exposure to DDT does not induce markers of GR disruption

While we found evidence of GR disruption in acute exposures to EDCs, we next evaluated the effect of chronic exposure to RU486 and DDT. Mice were provided with RU486 or DDT through their drinking water for 2 weeks or 1 month, then were tested for systemic markers of GR disruption. Throughout the study, mouse weights were monitored. We found that treatment with RU486 resulted in decreased weight, but this recovered by 1 month (Figure 4A). DDT did not affect weight. Circulating levels of corticosterone were tested using an ELISA. Treatment with RU486 for 1 month resulted in significantly increased corticosterone levels (Figure 4B). This aligns with previous studies using RU486 as a model [16-19], where glucocorticoid levels are increased with prolonged RU486 exposure. Next, we performed qRT-PCR on the adrenal glands to assess the expression of steroidogenesis genes. Neither RU486 nor DDT influenced *Star* or *Cyp11b1* expression (Figure 4C). Finally, gene expression of GR-responsive genes was evaluated in the liver. There were no significant changes to *Sgk1* expression, but RU486 treatment for 2 weeks significantly reduced *Tsc22c3* expression. These results are variable, with some alterations to body weight, GR transactivation in the liver, and corticosterone levels in RU486-treated mice. However, DDT did not significantly affect any of the tested markers of GR disruption.

**Figure 4.**
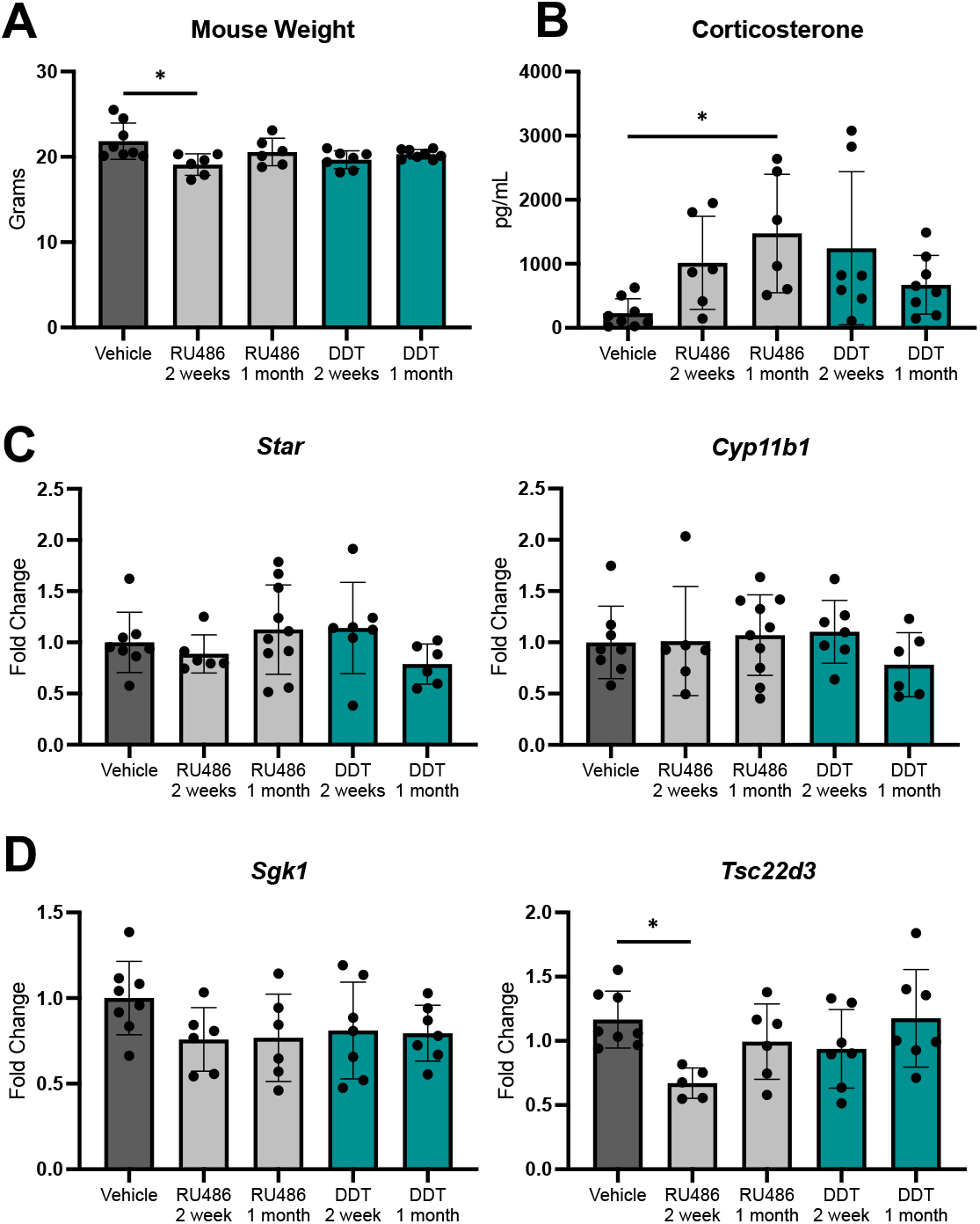
Chronic exposure to DDT did not elicit markers of GR disruption. C57BL/6 mice were exposed to RU486 or DDT through their drinking water for 2 weeks or 1 month. (A) Mouse weight at each time point. (B) Circulating levels of corticosterone. (C) qRT-PCR of steroidogenesis genes from RNA isolated in the adrenal glands. (D) qRT-PCR of GR-responsive genes from RNA isolatied from the liver. n≥6. *p≤0.05, **p≤0.01, ***p≤0.001, and ****p≤0.0001.

Glucocorticoids are known to have immunomodulatory and immunosuppressive effects [2]. To test the impact of chronic exposure to EDCs on immune cell populations, we performed flow cytometry on blood from mice exposed mice to RU486 or DDT for 1 month. We analyzed the lymphocyte populations, looking at B cells, T cells, and T cell subsets (Figure 5A). The overall immune cell counts were trending to be increased in mice treated with RU486 (Figure 5B). There was a significant reduction in B cells in both RU486 and DDT-treated mice (Figure 5C). T cells were significantly increased by RU486 treatment, but there was no difference in CD4+ and CD8+ T cell subsets (Figure 5D). The moderate effects of EDC treatment, both to systemic markers of disruption and immune cell populations, suggest a resiliency of the HPA axis, adapting to external changes to maintain homeostasis.

**Figure 5.**
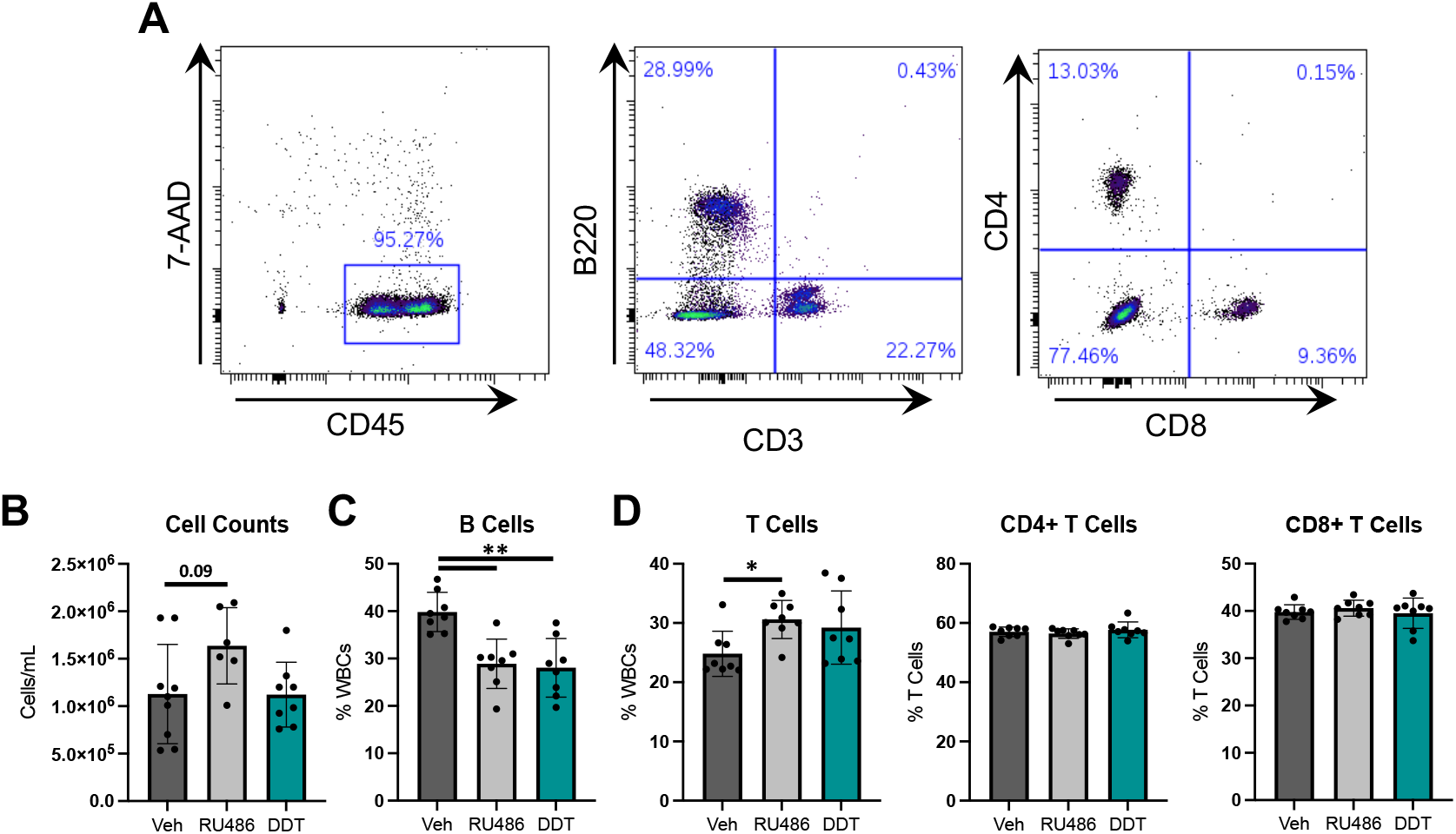
Chronic glucocorticoid receptor disruption suppresses circulating B lymphocytes. C57BL/6 mice were exposed to RU486 or DDT through their drinking water for 1 month. Flow cytometry was done on blood samples. (A) Gating strategy of flow cytometry analysis. (B) Total cell counts after isolation of white blood cells. (C) B cells and (D) T cells as a percent of CD45+ cells. n≥6. *p≤0.05, **p≤0.01, ***p≤0.001, and ****p≤0.0001.

## Discussion

Glucocorticoids are steroid hormones produced by the adrenal glands in a circadian manner and in response to stress. They are essential for life, regulating metabolism, immune function, cardiovascular function, and cognition [20]. Impaired regulation and signaling of glucocorticoids are associated with metabolic and cardiovascular disease, osteoporosis, immune dysfunction, mood disorders, and cancer [21]. These critical physiological processes mediated by glucocorticoids may be disrupted by exposure to environmental contaminants, increasing disease susceptibility. Identifying EDCs that can interfere with glucocorticoid signaling will provide crucial insights into several diseases and disorders. While several putative EDCs have been identified in vitro, few studies have evaluated their in vivo effects. In this study, we identified the agricultural agents DDT and ziram as GR disruptors *in vitro*. These chemicals influenced GR binding and translocation, leading to alterations in GR transactivation. Given the persistent presence of EDCs in the environment, chronic exposure may progressively alter GR function, contributing to an increased risk of inflammatory and metabolic diseases, particularly in populations with prolonged exposure.

The physiological effects of glucocorticoids are mediated by the glucocorticoid receptor (GR), a ligand-dependent transcription factor [22]. Under normal conditions, glucocorticoids bind to the GR, causing a conformation change. This allows the GR to translocate into the nucleus to bind glucocorticoid response elements (GREs) and initiate transcriptional changes. We evaluated DDT, metolachlor, and ziram for their binding ability to the GR and discovered DDT and ziram act as GR antagonists, while metolachlor did not bind GR (Figure 2A). Visualization of GR nuclear translocation revealed two distinct mechanisms by which EDCs impact glucocorticoid signaling. DDT induced nuclear translocation in the absence of cortisol, while ziram maintained the GR in the cytoplasm in the presence of cortisol (Figure 2B). These GR binding and translocation alterations resulted in transactivation disruption, decreasing the expression of *FKBP5* and *PER1* (Figure 1B). These findings highlight how exposure to EDCs can reduce GR-mediated gene expression, which compromises glucocorticoid function. This disruption may contribute to poor stress responses, heightened inflammation, and metabolic dysfunction, including insulin resistance and obesity, ultimately influencing long-term health outcomes.

A host of EDCs have been identified for the estrogen and androgen receptors, and their in vivo effects have been well described [23, 24]. In contrast, relatively few EDCs have been found to modulate GR activity, and even fewer have been validated in vivo [25]. Studies on the in vivo effects of GR disruption in mice present unique challenges. Unlike humans, where impaired glucocorticoid signaling is fatal without treatment, pathogen-free mice can readily survive complete loss of circulating glucocorticoids via adrenalectomy [26-28]. Thus, the physiological impact of GR disruption in mice during steady-state conditions may not be readily apparent.

Additionally, no standardized experimental guidelines exist for determining whether a chemical functions as a GR disruptor in vivo. In this study, we found that even the well-established GR antagonist RU-486 (mifepristone) caused only subtle GR disruption, leading to mild alterations in circulating corticosterone levels and modest changes in hepatic gene expression. These findings suggest that the hypothalamic-pituitary-adrenal (HPA) axis is highly resilient to external modulation by xenobiotics. However, in adrenalectomized (ADX) mice, treatment with RU-486 or DDT effectively suppressed the induction of GR target genes in the liver. These results indicate that while a corticosterone-challenge approach provides a useful initial framework for in vivo testing of potential GR-disrupting EDCs, standardized long-term assays are needed to fully assess their impact.

We showed here that chronic exposure to GR-EDCs suppressed the number of circulating lymphocytes. Interestingly, our results are the opposite compared to genetic depletion of the GR from the B cell lineage [29], suggesting either different mechanisms at play or that DDT caused B cell suppression independent of GR signaling. DDT has previously been shown to alter immune responses. Human PBMCs treated with 2.5µM DDT had increased IL-1B secretion [30]. Alternatively, an environmental study found that mice collected from DDT abatement sites had significantly lower leukocyte counts [31], supporting our findings that GR disruptors lead to less circulating lymphocytes. DDT has also been evaluated for its role in prenatal development. Prenatal exposure to DDT in rats leads to impaired morphogenesis of the adrenal cortex, reducing corticosterone and sex hormone production [32]. In a study analyzing cord blood transcriptomics, there was an increase in obesity-related transcripts in individuals with high levels of p,p’-DDE, a metabolite of DDT, suggesting an early role for EDC exposure in the risk of obesity and metabolic diseases [33]. These findings all implicate DDT as a key toxin, causing impaired immune function and increasing the risk of metabolic disease. However, more in vivo studies are needed to determine the role of DDT exposure on GR disruption.

The incidence of hypersensitivity, chronic inflammatory diseases, and autoimmune syndromes are on the rise in developed countries [34, 35]. The immune-suppressive effects of glucocorticoids are well-known, and disruption of glucocorticoid signaling has been shown to induce inflammation in a variety of organ systems including the stomach, small intestines, and heart [28, 36-38], promote autoimmune disease development [39, 40], and increase susceptibility to death by sepsis [41, 42]. Beyond their critical anti-inflammatory functions, emerging evidence suggests that endogenous glucocorticoids play a key role in maintaining immune system readiness. Recent studies indicate that impaired glucocorticoid signaling in T cells weakens antigen recognition, preventing effective T cell responses [43]. Furthermore, we recently reported that GR disruption in macrophages compromised the gastric immune response to *Helicobacter pylori* infection [44]. These findings highlight the potential consequences of exposure to GR-disrupting EDCs, which could significantly impact immune function and disease susceptibility [45]. However, studying the in vivo effects of GR-acting EDCs presents substantial challenges. As human exposure to these compounds continues to increase [46], it is imperative to conduct further research to clarify their impact on GR signaling, immune regulation, and inflammatory disease risk.

## Abbreviations

(HPA axis): Hypothalamic-Pituitary-Adrenal Axis
(GR): Glucocorticoid Receptor
(GREs): Glucocorticoid Response Elements
(EDCs): Endocrine-Disrupting Compounds
(DDT): Dichlorodiphenyltrichloroethane

---

**Supplementary Table 1.** Putative EDCs screened by a GR-responsive luciferase assay.

| Chemical | Manufacturer |
| --- | --- |
| Endrin | Sigma-Aldrich |
| Fipronil | Sigma-Aldrich |
| D-(+)-Cellobiose | Sigma-Aldrich |
| Monocrotophos | Sigma-Aldrich |
| Erythritol | Sigma-Aldrich |
| Diphenylamine | Sigma-Aldrich |
| 7-amino-4-methylcoumarin | Sigma-Aldrich |
| Diphenylmethane | Sigma-Aldrich |
| Resveratrol | Sigma-Aldrich |
| Vinclozolin | Cayman Chemicals |
| Bisphenol A | Sigma-Aldrich |
| Benomyl | Sigma-Aldrich |
| Ethiofencarb | Sigma-Aldrich |
| Anisole | Sigma-Aldrich |
| Parathion | Sigma-Aldrich |
| o-p'-DDT | Chem Service Inc |
| Metolachlor | Sigma-Aldrich |
| Ziram | Sigma-Aldrich |

**Supplementary Figure 1.**
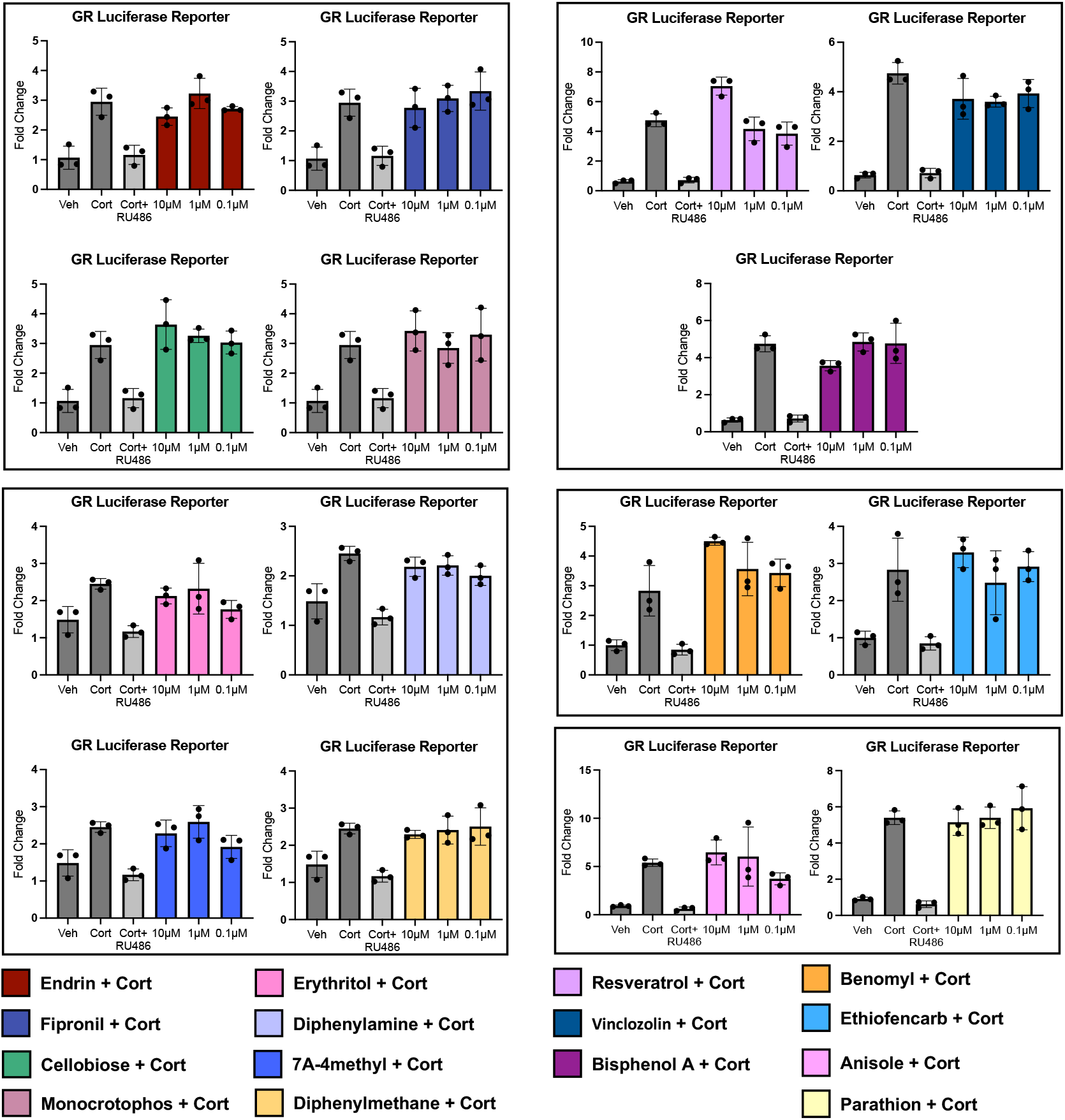
Putative EDCs that did not disrupt cortisol-induced luciferase expression. (A) A549 cells were transfected with a glucocorticoid-responsive luciferase construct. Cells were pretreated with RU486 or the putative EDCs for 1 hour, and then cortisol was added for 18 hours. Grouped results are from the same experiment and utilize the same controls. n=3. *p≤0.05, **p≤0.01, ***p≤0.001, and ****p≤0.0001.

## Notes

**Funding:** This work was supported by West Virginia University start-up funds (J.T.B). The West Virginia University Microscope Imaging Facility and Flow Cytometry & Single Cell Core receive support from the National Institutes of Health grants P30GM103503 and S10 grant OD028605, and U54 GM104942 respectively.

**Disclosures:** The authors have declared that no conflict of interest exists.

### Competing Interest Statement

The authors have declared no competing interest.

